# Zero-shot animal behavior classification with vision-language foundation models

**DOI:** 10.1101/2024.04.05.588078

**Authors:** Gaspard Dussert, Vincent Miele, Colin Van Reeth, Anne Delestrade, Stéphane Dray, Simon Chamaillé-Jammes

## Abstract

1. Understanding the behavior of animals in their natural habitats is critical to ecology and conservation. Camera traps are a powerful tool to collect such data with minimal disturbance. They however produce very a large quantity of images, which can make human-based annotation cumbersome or even impossible. While automated species identification with artificial intelligence has made impressive progress, automatic classification of animal behaviors in camera trap images remains a developing field.

2. Here, we explore the potential of foundation models, specifically Vision Language Models (VLMs), to perform this task without the need to first train a model, which would require some level of human-based annotation. Using an original dataset of alpine fauna with behaviors annotated by participatory science, we investigate the zero-shot capabilities of different kind of recent VLMs to predict behaviors and estimate behavior-specific diel activity patterns in three ungulate species.

3. Our results show that using these methods, it is possible to achieve accuracies over 91% in behavior classification and produce activity patterns that closely align with those derived from participatory science data (overlap indexes between 84% and 90%).

4. These findings demonstrate the potential of foundation models and vision-language models in ecological research. Ecologists are encouraged to adopt these new methods and leverage their full capabilities to facilitate ecological studies.

## 1 Introduction

Camera-traps are a powerful tool for studying wildlife in its natural habitats with minimal disturbance. As they are increasingly deployed throughout the world, a growing need for efficient methods to automatically analyze the vast amount of images they generate emerges. This is for instance true for one common goal of camera-trapping, which is to describe the diel activity pattern and time budget of some target species, i.e. the distribution of their level of activity and behaviors throughout the day (J. M. Rowcliffe et al. 2014; Gilbert et al. 2023).

Deriving activity patterns from camera-traps images first requires identifying the species present in the pictures (Whytock et al. 2021). This task can now often be fully automatized, as recent machine learning models achieve very high performance in species classification (Rigoudy et al. 2023; Willi et al. 2019; Mitterwallner et al. 2023). Then, activity level is generally assumed to be proportional to the number of sightings. One simply counts the number of sightings of the species of interest over time intervals (e.g. hours) and build activity curves (Ridout and Linkie 2009), or make more complex modelling of the distribution of sightings (Gerber et al. 2024). However, one might want to study more informative, behavior-specific patterns, for instance to understand when are animals foraging, resting or moving. In contrast with species identification, classification of animals’ behavior in camera-trap images or video is still in its infancy (Norouzzadeh et al. 2018; Liu et al. 2023; Schindler and Steinhage 2021; Schindler, Steinhage, et al. 2024).

In addition to identifying the best models for this task, several key constraints for developing such models exists: firstly, a large amount of training data is needed, but obtaining expert labels of behaviors on large sets of images is very time-consuming and costly. Behaviorally-annotated camera-trap datasets are currently very rare; secondly, a lot of computational resources are required for training. Such resources might not be available to ecologists, and training has a large carbon footprint; thirdly, the model need to be re-trained each time a new species or behavior class is added.

However, research groups in artificial intelligence worldwide are publishing “foundation models”, capable of executing zero-shot tasks, i.e. capable of making predictions on tasks for which they have not been explicitly trained (Radford et al. 2021; Kirillov et al. 2023). These pre-trained foundation models can solve most of the above-mentioned problems, since they don’t require training data, inference requires much less energy than training and the billions of examples seen during training theoretically allow them to predict and generalize on various species and behavior of interest (Fabian et al. 2023).

In this article, we investigate whether foundation models are already performant enough to be used in ecology for animals’ behavioral classification, from camera-trap images, without any fine-tuning. We use an original dataset of camera trap images of European fauna with behavior labels provided by participatory sciences. We explore the zero-shot capabilities of six recent Vision Language Models (VLM)(Bordes et al. 2024) to predict the behavior of three different species. As it is known that the formulation of the prompt request has a significant impact on the output of the models, we explore the importance of this effect for the proposed task. In particular, as we hypothesize that the models have not been trained on many images of wild animals active at night and may be biased towards certain behaviors. We therefore also investigate the effect of specifying to the model whether the image was taken during nighttime or daytime. In addition to accuracy measures, we use overlap measures to quantify to what extent diel activity patterns, constructed independently for each behavior and from human annotations, can be well estimated by relying only on models’ predictions. Finally, we discuss how our study demonstrates the power of vision-language models that can immediately assist ecologists in many cumbersome tasks.

## 2 Material and Methods

### 2.1 Dataset

The CREA Mont-Blanc camera trap dataset is collected by CREA Mont-Blanc (Centre de Recherches sur les Ecosystèmes d’Altitude) in the valley of Chamonix in the French Alps. The dataset is made of 131 537 images collected between 2017 and 2020 during 24 580 events, events being defined as 1-minute periods. Images were labelled on https://www.zooniverse.org/ using participatory science. Participants were asked to identify the species and could also vote for the animal’s behavior using five non-exclusive classes: *eating*, *resting*, *moving*, *standing* and *interacting*. Each event had been viewed by at least 30 people at the end of the process. The distribution of species and behaviors is shown in figure Figure 1a. We have focused our work on three species that are actively studied by the CREA Mont-Blanc: chamois (*Rupicapra rupicapra*), red deer (*Cervus elaphus*) and roe deer (*Capreolus capreolus*), and focused on 3 behaviors: eating, moving and resting. We excluded standing from our analysis due to inconsistent interpretation by participants and its frequent combination with other behaviors. Additionally, we did not focus on interacting since there were only 71 events with this behavior in the entire dataset.

**Figure 1:**
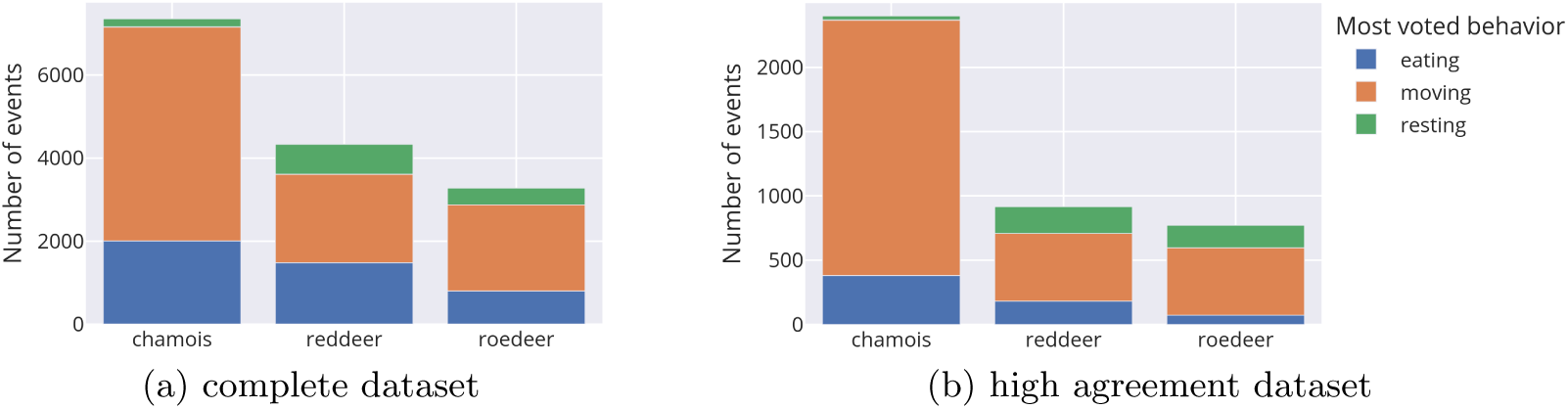
Species and behavior distribution of the CREA Mont-Blanc camera trap dataset for the three species of interest. (a) with all events, (b) with events for which the most voted behavior has at least 90% of the votes

### 2.2 Zero-shot Vision-Language Models

Vision-Language Models (VLMs) are multi-modal deep learning models that can process both image and text simultaneously. These models, often trained on very large datasets, demonstrate very good zero-shot performances in downstream tasks. Here, we investigate the zero-shot capabilities of two categories of VLMs: contrastive vision-language models and multimodal Large Language Models (LLMs). A brief overview of these two categories is given below, for a comprehensive review, see Bordes et al. (2024).

#### Contrastive Vision-Language Models

These models are trained to predict the similarity between a text and an image. Formally, a vision encoder and a text encoder are used to extract embeddings of images and texts, and the similarity of an image-text pair is obtained through the cosine similarity of the normalized embeddings (see Figure 3a). During the training, a contrastive image-text objective is used to bring closer the embeddings of positive image-text pairs and push apart those of negative pairs. In our study, we decided to use CLIP (Radford et al. 2021) for our baseline as the first contrastive vision-language model of large scale, and SigLIP (Zhai et al. 2023) which improved on CLIP with a new contrastive objective. We also include WildCLIP(Gabeff et al. 2023), a fine-tuned version of CLIP trained exclusively on camera trap images of African fauna. WildCLIP uses the large Snapshot Serengeti dataset (Swanson et al. 2015), with image captions built from attributes labeled through participatory science, including animal behavior.

#### Multimodal Large Language Models

These models process both image and text tokens at the input of a Large-Language Model (LLM). The image tokens are obtained using a vision encoder, typically a pretrained contrastive model such as CLIP. The integration of a language model allows these models to perform more complex reasoning compared to contrastive models. We examine three multimodal LLMs of varying sizes: CogVLM (Wang et al. 2024), which has a very large vision encoder (EVA2-CLIP-E); MobileVLM V2 (Chu et al. 2024), a lightweight VLM which uses CLIP ViT-L/14; and the more recent PaliGemma (Beyer et al. 2024), which utilizes SigLIP-SO.

### 2.3 Zero-shot inference

For all methods, images are first cropped using MegaDetector v5 (Beery et al. 2019) to ensure that the image encoder of each model extracts information related to the animal and not to the rest of the scene.

Contrastive vision-language models such as CLIP are able to perform zero-shot classification (Radford et al. 2021) using their ability to predict the similarity score between an image and a text. Here, we follow this methodology and match the images with the following captions: “a camera trap picture of an animal *{*behavior*}*”. This process is illustrated on Figure 3a. We calculate similarity scores for each behavior, normalize them using the softmax function, and select the behavior with the highest score as the final prediction. We use the normalized similarity score of the chosen behavior as the confidence score.

For the multimodal LLMs, we perform a Visual Question Answering task (Agrawal et al. 2016). As shown on Figure 3b, we prompt the assistant with the following question: “Is the animal in the image eating, moving or resting? Answer the question using a single word.”. For each word prediction in the answer, the model considers all possible words in the vocabulary, assigns a score to each one and predicts the one with the highest score. In our case, since the model predicts directly the behavior with the first word, we can use it to retrieve the score of each behavior. Then, following the methodology described by Jiang et al. (2021), the score of the predicted behavior is normalized by the sum of the scores of the candidates words only (the three behaviors words).

We also experiment to give additional context to the models to improve their performances: we inform the model whether the image was taken during daytime or nighttime. For CLIP-based methods, we change the caption to “a camera trap picture of an animal *{*behavior*}*, captured during *{*context*}*.” with *context* being either “nighttime” or “daytime”. For LLMs, we add “This image is captured during *{*context*}*.” before the question.

For each behavior, the classification performance of the model are evaluated with the accuracy and the calibration of the scores is measured with the ECE (Guo et al. 2017). For these two metrics, we use a subset of the dataset that removes ambiguity in the labels (e.g. sequences with an animal that is seen both eating and moving). This subset is obtained by keeping only the images for which the most voted for behavior has at least 90% of the votes, and will be called the *high agreement* dataset hereafter. To address the imbalance between behaviors (Figure 1b), we report the macro accuracy in addition to the accuracy of each behavior. This metric is calculated by taking the average of the accuracy of the three behaviors, each given equal weight, ensuring that no single behavior disproportionately influences the overall accuracy. We also aggregate individual images predictions to event predictions to improve the accuracy of the models (Dussert et al. 2024). Confidence scores at the sequence level are calculated by averaging the scores of the images in the sequence.

### 2.4 Behavior-specific activity patterns

Diel activity patterns are computed either on the Zooniverse annotations or on the models predictions using the time at which the camera trap events were triggered. As proposed by Vazquez et al. (2019), times are first converted to radian and standardized using two anchor time (the average sunrise time and the average sunset time) to account for the variation of the length of the day during the year. A circular kernel density (of the von Mises distribution) is fitted to the radian times to obtain the activity pattern. The overlap index Δ is used to compare two activity patterns by computing the integral of the minimum between the two density function (Ridout and Linkie 2009). An index of 1 is obtained for a perfect overlap and 0 for no overlap at all. In practice, it is computed with the estimator 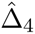 of Ridout and Linkie (2009). These operations were done with the functions *solartime*, *fitact* and *compareCkern* of the activity package in R (M. Rowcliffe 2023), with 200 bootstrap replicates for the estimation of 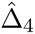.

Activity patterns are computed using the complete dataset for our three species and three behaviors of interest. For Zooniverse annotations, we keep only the events for which a behavior has at least 50% of the votes. For models prediction, we used the same threshold of 0.75 on the confidence scores of all the models to reduce false positives. We did not apply any filter to ensure independence between event (e.g. 30 minutes interval between two events) as it has been shown to bias the activity patterns (Peral et al. 2022).

## 3 Results

Starting with the contrastive vision-language models, the baseline CLIP with ViT-Base architecture performs poorly for this behavior recognition task, with a low accuracy of 71.75% (Table 1). It particularly fails at correctly predicting the eating behavior. The SigLIP model represents a significant improvement over the CLIP model, with a much better accuracy of 91.14%. The accuracy of the eating behavior is however still lower than the one for the other behaviors (Table 1). The specialized WildCLIP model offers very similar performance to SigLIP despite a much smaller number of parameters. For the multimodal LLMs, CogVLM gives the best performance, with an accuracy of 97.45%. PaliGemma which uses SigLIP as vision encoder is only slightly better than SigLIP but performs better than MobileVLM V2 despite having less parameters. Per-species accuracies are shown in Supporting Information Table 1.

**Table 1:** Behavioral classification accuracy and expected calibration error (ECE) of the six studied models on the high agreement dataset.

| Model | Context (day & night) | Accuracy |  |  | Macro accuracy | ECE (%) | #Params |
| --- | --- | --- | --- | --- | --- | --- | --- |
|  |  | Eating | Moving | Resting |  |  |  |
| Contrastive models |  |  |  |  |  |  |  |
| CLIP (ViT-B/16) | × | 31.76 | 92.39 | 91.08 | 71.75 | 17.15 | 150M |
|  | ✓ | 48.11 | 88.67 | 93.98 | 76.92 | 19.61 |  |
| SigLIP (ViT-SO400M) | × | 84.28 | 94.20 | 94.94 | 91.14 | 10.78 | 878M |
|  | ✓ | 94.65 | 89.20 | 95.18 | 93.01 | 14.15 |  |
| WildCLIP (ViT-B/16) | × | 89.47 | **94.70** | 93.49 | 92.55 | **6.85** | 150M |
|  | ✓ | 90.09 | 94.17 | 95.42 | 93.23 | 7.80 |  |
| Multimodal LLMs |  |  |  |  |  |  |  |
| CogVLM | × | 99.37 | 93.71 | **99.28** | **97.45** | 11.93 | 18B |
|  | ✓ | **99.53** | 87.95 | 99.04 | 95.50 | 11.29 |  |
| MobileVLM V2 | × | 95.75 | 82.23 | **99.28** | 92.44 | 9.55 | 7B |
|  | ✓ | 96.07 | 78.89 | **99.28** | 91.41 | 10.35 |  |
| PaliGemma | × | 90.57 | 91.80 | 96.87 | 93.08 | 7.46 | 3B |
|  | ✓ | 92.30 | 91.41 | 95.42 | 93.04 | 9.38 |  |

Giving additional day-night context improves the accuracy of all the contrastive models (Table 1), but decreases the accuracy of the LLMs. None of the models exhibited a good score calibration, with the best model (WildCLIP) having an ECE of 6.85%. Moreover, adding context slightly deteriorates the calibration of almost all the models.

As expected from the heterogeneity in the models’ accuracy, the quality of the behavior-specific diel activity patterns estimated from their predictions varies. Figure 4 presents the activity patterns estimated using CLIP, SigLIP and CogVLM models without day/night context. These models were chosen to provide a comprehensive comparison across different classification accuracy levels. For other models, see Supporting Information Figure 1-6 and Table 2. Activity patterns computed with CogVLM predictions are very close to those obtained with participatory science labels, and even small variations in activity along the course of the day are captured. The activity patterns predicted by CLIP are generally unreliable. In particular, the model is biased towards predicting moving at nighttime. To a lesser extent, SigLIP also presents a night bias, which is particularly visible using the red deer data (Figure 4). This bias seems to be considerably reduced by the addition of the day/night context (Supporting Information Table 2).

**Figure 2:**
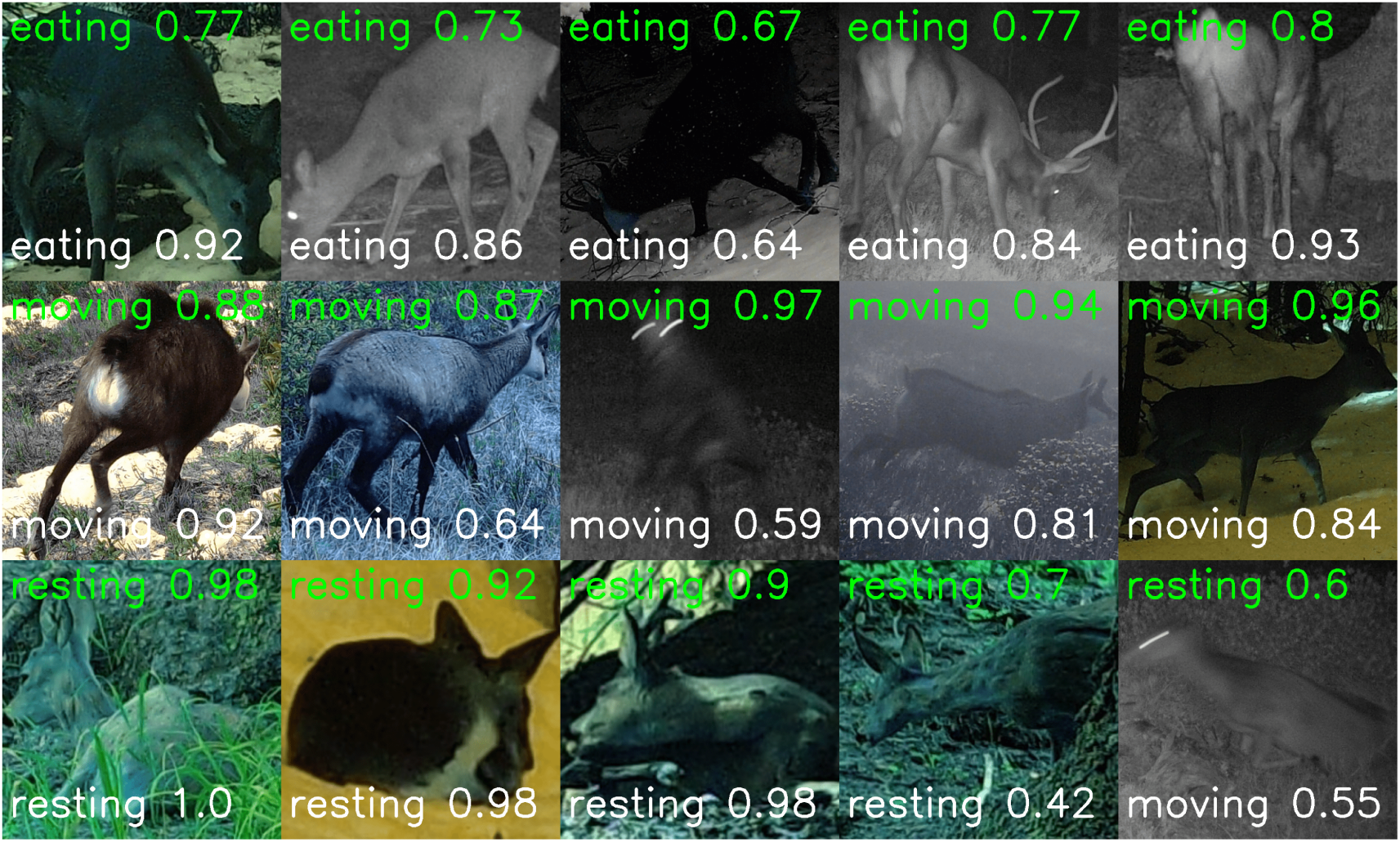
Sample images from the dataset, of the three species and three behaviors of interest. When an event has multiple image, only the first one is shown. The Zooniverse label and proportion of vote is shown at the top in green, and the prediction of the best zero-shot model (CogVLM) with its confidence score is shown at the bottom in white.

**Figure 3:**
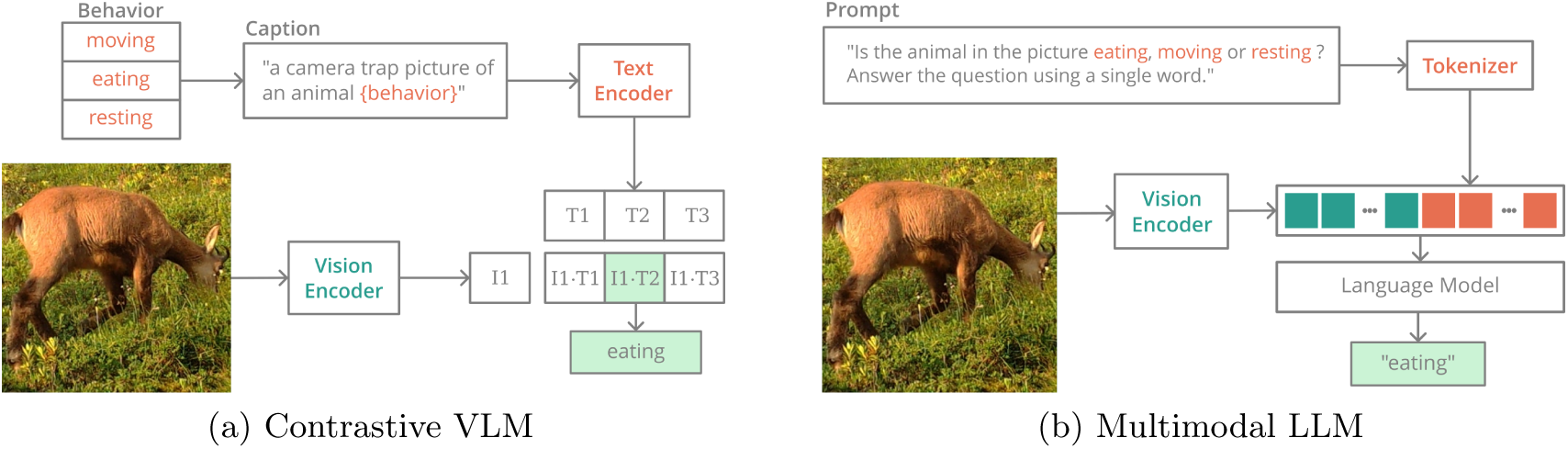
Overview of the zero-shot inference process for the two model families we tested. Contrastive vision-language model (a) predict the similarity between the image and the three captions, one for each behavior. The predicted behavior is the one with the highest similarity score (light blue). Multimodal LLMs (b) are language models that can process visual tokens in addition to the prompt, and predict the behavior as the next word in the sequence.

**Figure 4:**
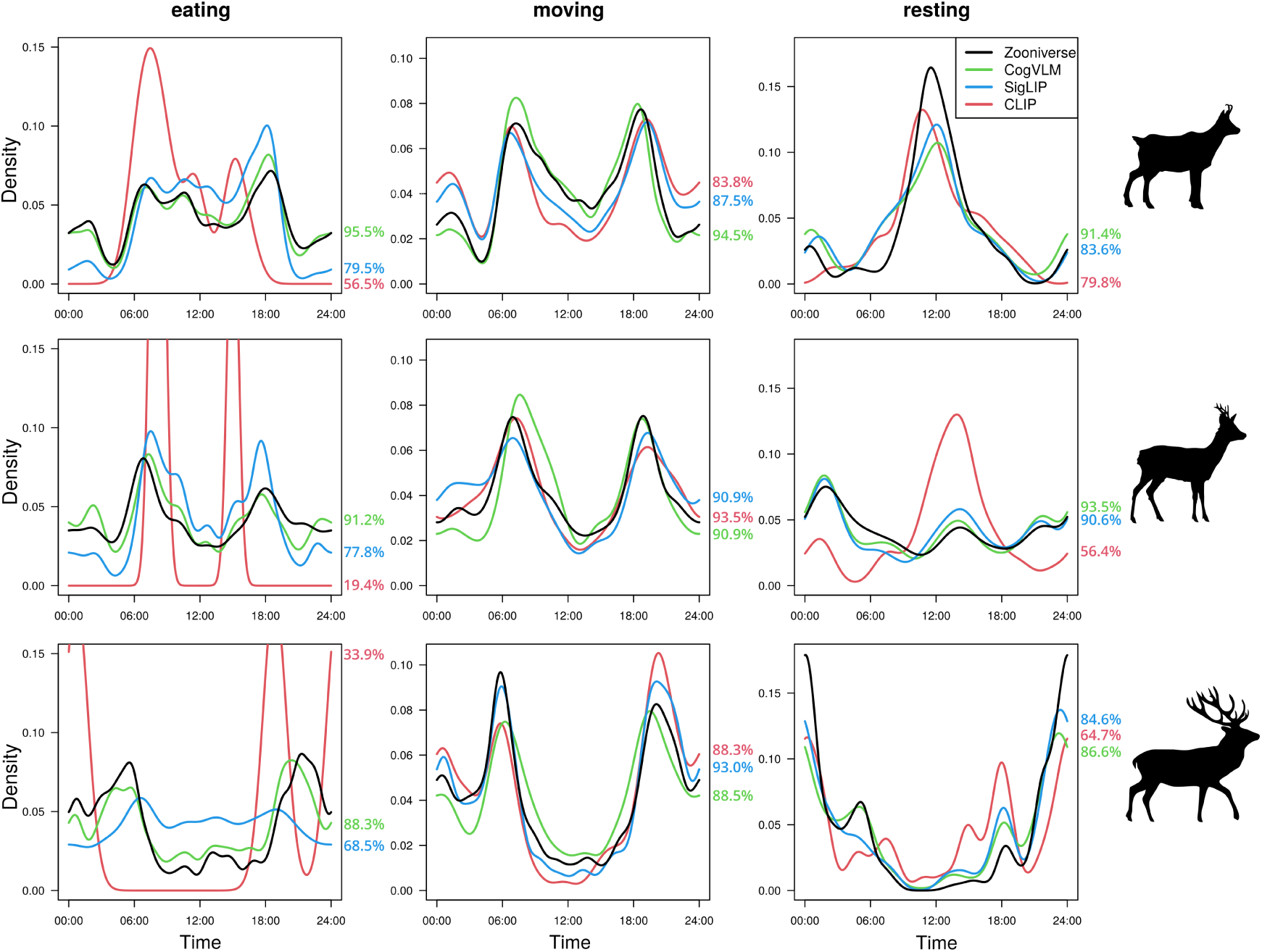
Comparison of the activity patterns using either the participatory science labels from Zooniverse (black) or predictions of three models (colors). Each row corresponds to a different species (top to bottom : chamois, roe deer, red deer). Overlap values are indicated on the right of the curves.

**Table 2:** Overlap index averaged over the three species of interest on the complete dataset.

| Model | Context | Average overlap index |  |  |  |
| --- | --- | --- | --- | --- | --- |
|  |  | Eating | Moving | Resting | All |
| Contrastive models |  |  |  |  |  |
| CLIP (ViT-B/16) | × | 36.60 | 66.97 | 88.53 | 64.03 |
|  | ✓ | 39.73 | 75.80 | 91.27 | 68.93 |
| SigLIP (ViT-SO400M) | × | 75.27 | 86.27 | 90.47 | 84.00 |
|  | ✓ | 89.97 | 85.13 | **93.73** | 89.61 |
| WildCLIP (ViT-B/16) | × | 91.03 | 80.63 | 92.27 | 87.98 |
|  | ✓ | 91.20 | 80.23 | 91.00 | 87.48 |
| Multimodal LLMs |  |  |  |  |  |
| CogVLM | × | 91.67 | 87.17 | 91.30 | **90.04** |
|  | ✓ | 93.00 | 87.37 | 85.27 | 88.54 |
| MobileVLM V2 | × | **93.43** | 83.37 | 87.67 | 88.16 |
|  | ✓ | 93.10 | 86.86 | 83.63 | 87.86 |
| PaliGemma | × | 88.17 | **90.47** | 82.83 | 87.15 |
|  | ✓ | 89.00 | 89.13 | 83.30 | 87.14 |

Quantitative measures of the overlap in activity patterns between the one derived from Zooniverse annotations and those derived from the various models are shown in Table 2. In accordance with previous results, CogVLM has the best overall overlap values, with an average overlap index of 90.04%. SigLIP, WildCLIP, PaliGemma and MobileVLM V2 exhibit slightly lower performance compared to CogVLM, while CLIP performs significantly worse. Once again, the additional day-night context helps with contrastive models, but not with LLMs. In particular, SigLIP with context emerges as the second-best model.

## 4 Discussion

This study evaluates the ability of zero-shot modern VLMs to predict the behavior of three mountain ungulates in camera trap images. We show that these models can have very good performances, with macro accuracy exceeding 91% for all models except the baseline (CLIP ViT-B). Furthermore, the diel activity patterns generated by these models closely align with those obtained through participatory science, exhibiting overlap indexes between 84% and 90%, again with the exception of CLIP.

Using an automated and flexible method, such as the one presented in this study, to classify animal behaviors on pictures can save considerable time compared to having images annotated by citizen scientists or experts. For instance, the very good SigLIP model can process tens of thousands of pictures in an hour (see below for further discussion). Therefore, this approach could enable large-scale annotation of camera trap datasets and facilitate or automate the study of behaviors, which could for example be particularly useful for studying the effects of climate change or human activities on the behavior of wild animals. These research topics are increasingly studied using camera-traps (Bison et al. 2024; Blount et al. 2024). Furthermore, even though this study illustrates the effectiveness of foundation vision-language models for a specific task, the flexibility of the method makes it easily adaptable to other ecological tasks. For example, it could be possible to predict human behaviors or landscapes attributes by changing the prompt (or the caption) given to the model.

While some existing datasets such as MammalNet (Chen et al. 2023) and AnimalKingdom (Ng et al. 2022) are available and have more species and behaviors, we decided not to use them as they consist of very high quality video footage. Good model performance on these datasets may not transfer to camera trap datasets given the distribution gap between the two. In addition, we decided not to focus on video processing, as many camera trap studies use single shot or burst mode, requiring the development of methods to obtain the most accurate behavior possible from one or a few images (Rovero and Zimmermann 2016).

A visual analysis of discrepancies between model predictions and participatory science annotations reveals a degree of inherent ambiguity and that both may be correct in certain instances. In fact, some sequences of pictures clearly contains images in which the behavior of the animal differ (e.g. Figure 2, bottom right), with eating and moving being the most frequent combination. While not common, these ambiguities may explain the slight deviation in the diel activity curves obtained from CogVLM, despite its near-perfect accuracy on the high agreement dataset. This suggests that exploring a zero-shot multilabel classification approach might be useful. Furthermore, the models occasionally make errors attributable to a poor scene understanding: an animal walking in the foreground might be misclassified as "eating" if its head aligns with background vegetation, or misclassified as "resting" if its legs are hidden by deep snow or rocks. These errors could potentially be eliminated using VLMs capable of simultaneously processing multiple images from the same sequence, rather than independently.

Given that several of the models achieve very good performance, it is interesting to study the trade-off between model size and performance. When considering multimodal LLMs, CogVLM appears to be the best model if inference time is not a concern while MobileVLM V2 is clearly not interesting, as its performance is similar to the one of WildCLIP, which is 50 times smaller. The difference in performance is highly likely due to the differing sizes of the vision encoders between CogVLM (10B parameters) and MobileVLM (428M parameters). For contrastive models, WildCLIP seems to be the most efficient model. It performs considerably better than CLIP ViT-Base, despite having the same number of parameters. WildCLIP’s performance is impressive given that it has been fine-tuned on African data only, which demonstrates the effectiveness of the VR-LwF "Learning without Forgetting" loss that was used. However, the model is not performing better than SigLIP despite having already been exposed to camera trap images and similar captions depicting the same behaviors. This may be attributed to WildCLIP having much fewer parameters and the distribution shift between the training data and our dataset. Nonetheless, the release of our data will facilitate the development of specialized models more robust to the changes in species, habitats and behaviors. Finally, SigLIP seems to be the most efficient non-specialized foundation model and has a reasonable inference time (1 hour for 40 000 images on one NVIDIA V100 GPU), which could also make it useful for analyses conducted on standard desktop CPU-only computers.

Providing the context of whether the image was taken during the day or at night only seems to improve the contrastive models. This can be explained by the fact that giving a more precise caption reduces the distribution gap with the captions in the training dataset (Radford et al. 2021). Current multimodal LLMs, on the other hand, are trained with rigid conversation templates and can be very sensitive to the structure of the prompt. Most of the datasets they were trained on do not include context before the question. Hence, in contrast to the contrastive models, giving this additional context has increased the distribution gap and may have made confused the models. Moreover, current LLMs are not invariant to permutations in the order of choices presented in the prompt (Zong et al. 2023). In other words, the sequence in which options are listed can impact the prediction of the model. For example, given choices A, B, and C, the model may output different predictions if the options are presented as A-B-C or C-A-B. This is for instance the case in our study (see Supporting Information Table 3), although it does not alter our general conclusions. LLMs may also exhibit positional bias, being more likely to predict one choice based on its position rather than its content (Zheng et al. 2024). Therefore, while multimodal LLMs demonstrate impressive performance, it is crucial to acknowledge and account for these limitations in their application and interpretation of results.

Overall, our study raises questions regarding the necessity of training new models for ecological applications, especially considering the rapidly accelerating rate of publication of foundation models. We can expect significant improvements and capabilities in the near future, such as the ability to analyze multiple consecutive images, as the commercial GPT4-Vision model already does (OpenAI 2023). Ecologists should embrace these new methods, evaluate their performance, and learn how to exploit them to their full potential.

## Acknowledgements

This work was granted access to the HPC resources of IDRIS under the allocation 2022-AD010113729 made by GENCI.

## Author contributions

G.D., S.C.J., S.D. and V.M. conceived the ideas and designed the methodology. A.D. and C.V.R gathered, annotated and processed the CREA-Mont Blanc dataset. G.D. coded and performed the analysis. G.D. wrote the first version of the manuscript, S.C.J., S.D., A.D and C.V.R and V.M. contributed critically to the drafts and gave final approval for publication.

## Data availability statements

The code and dataset will be made available on Zenodo: https://doi.org/10.5281/zenodo.10925926

## Supporting information

**Table 3:** Behavior classification accuracy of the six models, for each species and behavior, using the high agreement dataset.

| Model | Context | Chamois |  |  | Roe deer |  |  | Red deer |  |  |
| --- | --- | --- | --- | --- | --- | --- | --- | --- | --- | --- |
|  |  | Eating | Moving | Resting | Eating | Moving | Resting | Eating | Moving | Resting |
| Contrastive models |  |  |  |  |  |  |  |  |  |  |
| CLIP | × | 91.65 | 36.48 | 93.75 | 93.51 | 26.03 | 89.14 | 94.11 | 24.18 | 92.31 |
|  | ✓ | 87.72 | 54.59 | 87.50 | 90.65 | 43.84 | 93.71 | 90.30 | 36.26 | 95.19 |
| SigLIP | × | 95.22 | 85.04 | 96.88 | 91.60 | 89.04 | 94.86 | 92.97 | 80.77 | 94.71 |
|  | ✓ | 92.65 | 94.49 | 93.75 | 80.53 | 97.26 | 95.43 | 84.79 | 93.96 | 95.19 |
| WildCLIP | × | 96.48 | 88.19 | 78.12 | 90.27 | 87.67 | 93.71 | 92.40 | 92.86 | 95.67 |
|  | ✓ | 95.87 | 89.24 | 81.25 | 90.84 | 87.67 | 96.00 | 91.06 | 92.86 | 97.12 |
| Multimodal LLMs |  |  |  |  |  |  |  |  |  |  |
| CogVLM | × | 95.87 | 99.21 | 100.0 | 91.03 | 100.0 | 99.43 | 88.21 | 99.45 | 99.04 |
|  | ✓ | 92.35 | 99.21 | 100.0 | 80.34 | 100.0 | 98.86 | 78.90 | 100.0 | 99.04 |
| MobileVLM V2 | × | 85.41 | 94.75 | 100.0 | 77.48 | 100.0 | 99.43 | 75.29 | 96.15 | 99.04 |
|  | ✓ | 82.64 | 94.75 | 100.0 | 72.71 | 100.0 | 99.43 | 70.91 | 97.25 | 99.04 |
| PaliGemma | × | 92.50 | 90.29 | 96.88 | 90.84 | 90.41 | 94.86 | 90.11 | 91.21 | 98.56 |
|  | ✓ | 92.60 | 91.86 | 96.88 | 88.93 | 93.15 | 93.14 | 89.35 | 92.86 | 97.12 |

**Table 4:** Overlap index of the six models, for each species and behavior, using the complete dataset.

| Model | Context | Chamois |  |  | Roe deer |  |  | Red deer |  |  |
| --- | --- | --- | --- | --- | --- | --- | --- | --- | --- | --- |
|  |  | Eating | Moving | Resting | Eating | Moving | Resting | Eating | Moving | Resting |
| Contrastive models |  |  |  |  |  |  |  |  |  |  |
| CLIP | × | 56.5 | 83.8 | 79.8 | 19.4 | 93.5 | 56.4 | 33.9 | 88.3 | 64.7 |
|  | ✓ | 65.8 | 90.4 | 77.2 | 18.2 | 92.4 | 72.7 | 35.2 | 91.0 | 77.5 |
| SigLIP | × | 79.5 | 87.5 | 83.6 | 77.8 | 90.9 | 90.6 | 68.5 | 93.0 | 84.6 |
|  | ✓ | 92.0 | 96.8 | 83.4 | 92.2 | 92.6 | 88.1 | 85.7 | 91.8 | 83.9 |
| WildCLIP | × | 93.0 | 97.0 | 71.8 | 88.9 | 90.7 | 85.6 | 91.2 | 89.1 | 84.5 |
|  | ✓ | 93.3 | 97.3 | 68.7 | 89.9 | 88.3 | 86.6 | 90.4 | 87.4 | 85.4 |
| Multimodal LLMs |  |  |  |  |  |  |  |  |  |  |
| CogVLM | × | 95.5 | 94.5 | 81.4 | 91.2 | 90.9 | 93.5 | 88.3 | 88.5 | 86.6 |
|  | ✓ | 96.5 | 89.3 | 81.6 | 92.9 | 83.2 | 93.9 | 89.6 | 83.3 | 86.6 |
| MobileVLM V2 | × | 95.9 | 94.4 | 73.8 | 91.6 | 84.6 | 92.4 | 92.8 | 84.0 | 83.9 |
|  | ✓ | 95.9 | 92.7 | 74.6 | 91.0 | 82.3 | 92.8 | 92.4 | 85.5 | 83.5 |
| PaliGemma | × | 89.9 | 94.1 | 71.8 | 90.5 | 89.4 | 88.9 | 84.1 | 87.9 | 87.8 |
|  | ✓ | 90.4 | 93.7 | 75.1 | 91.6 | 87.0 | 87.8 | 85.0 | 86.7 | 87.0 |

**Figure 5:**
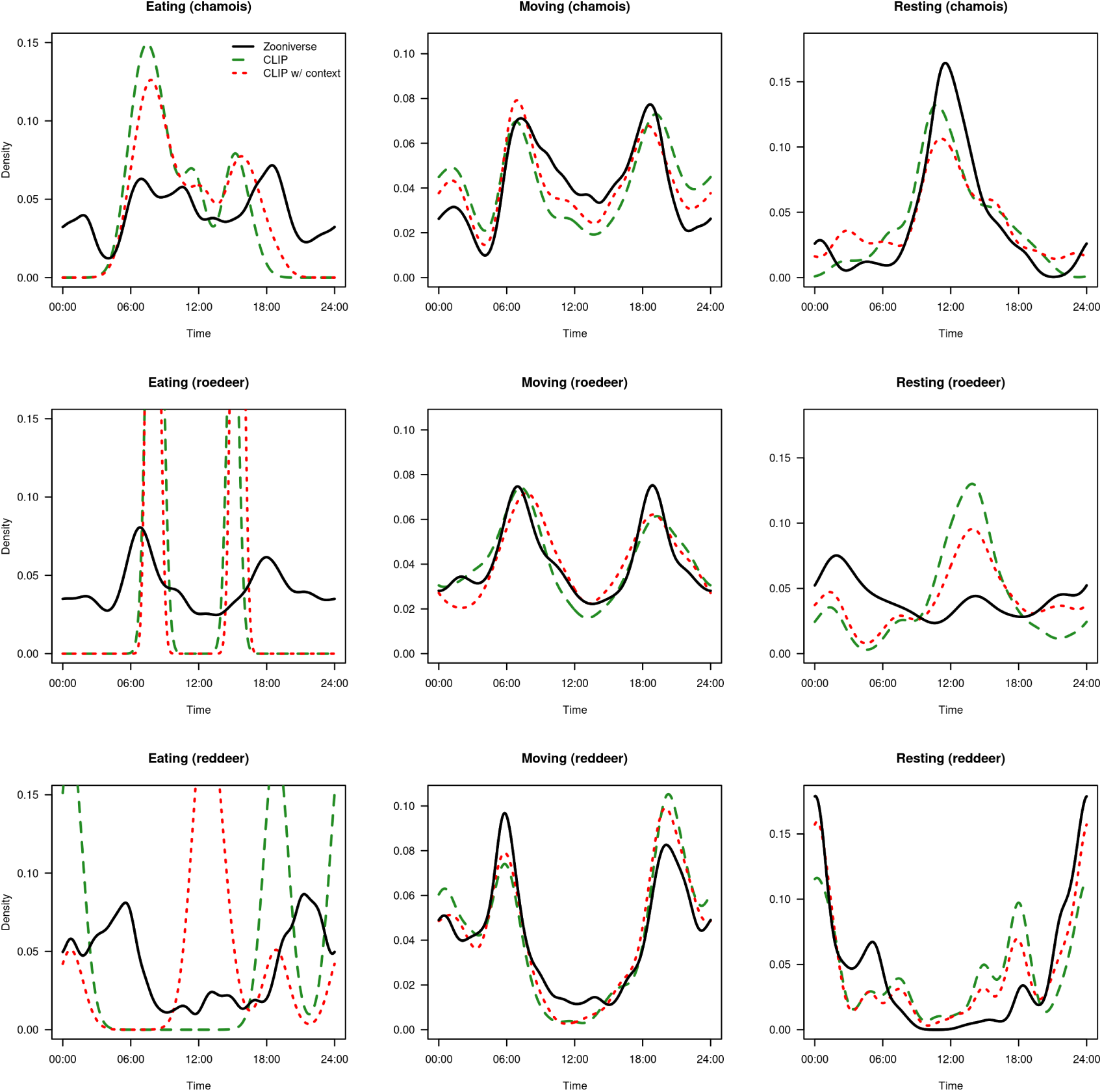
Activity patterns of CLIP, with and without day-night context, using the complete dataset.

**Figure 6:**
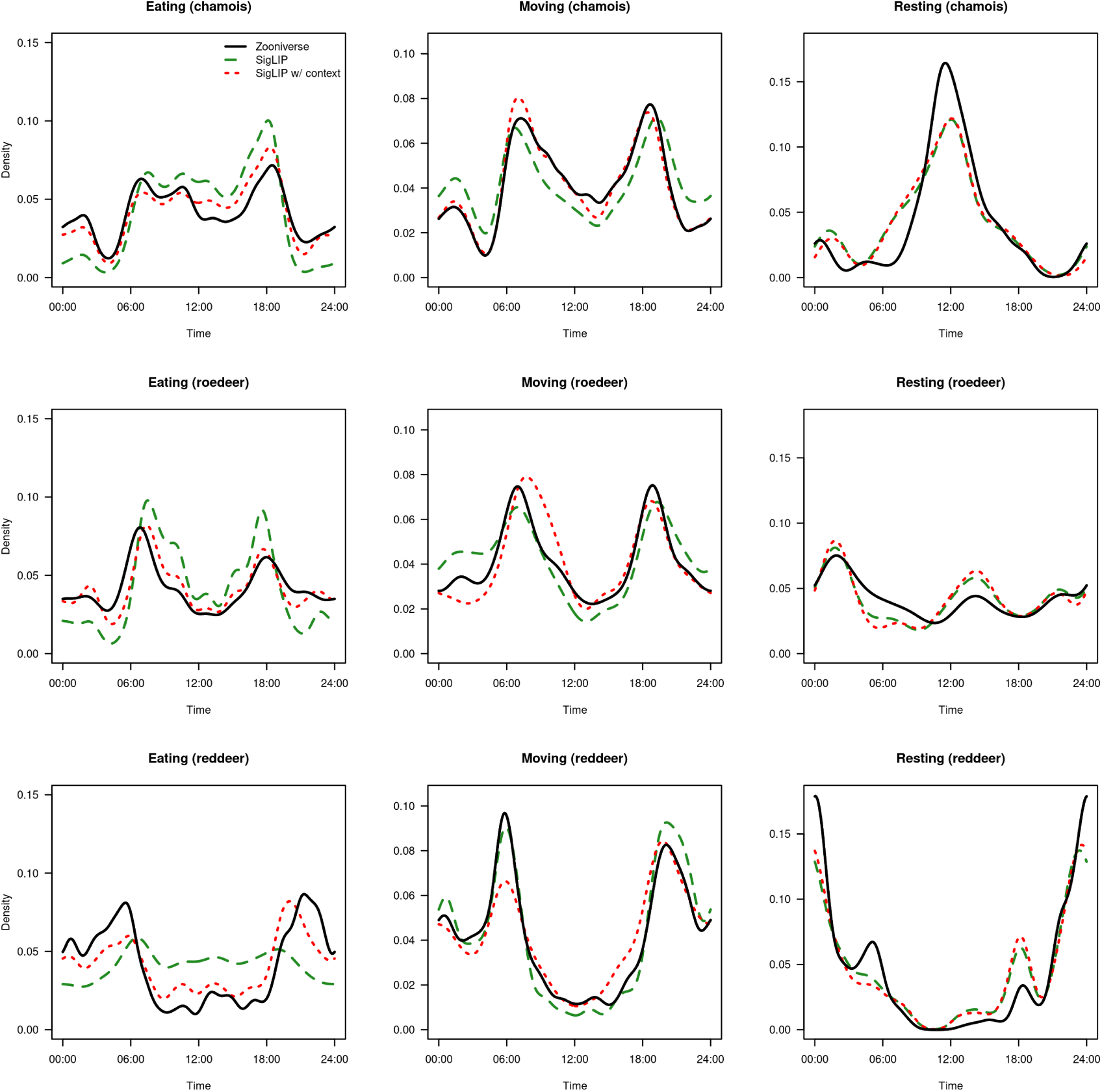
Activity patterns of SigLIP, with and without day-night context, using the complete dataset.

**Figure 7:**
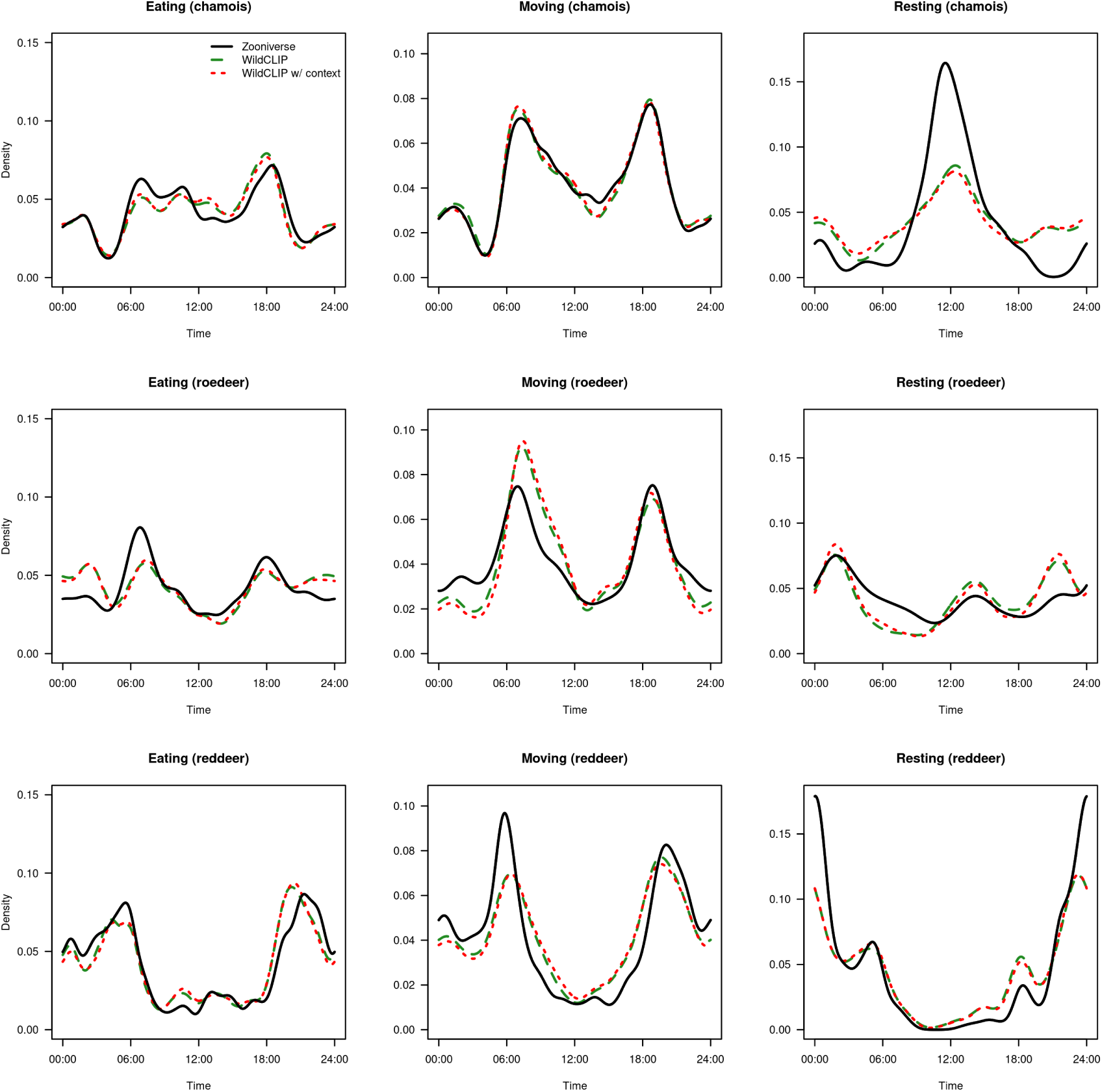
Activity patterns of WildCLIP, with and without day-night context, using the complete dataset.

**Figure 8:**
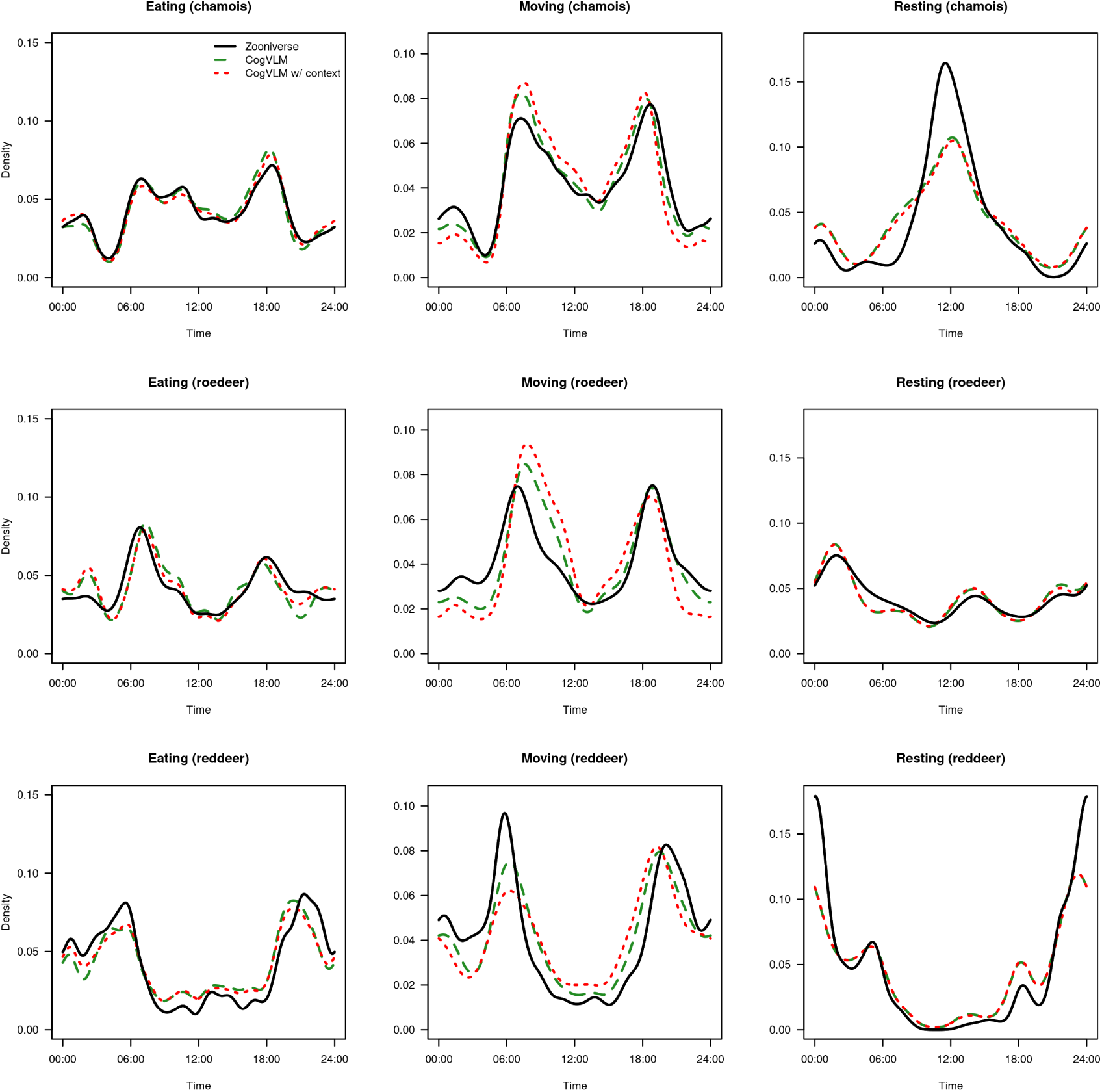
Activity patterns of CogVLM, with and without day-night context, using the complete dataset.

**Figure 9:**
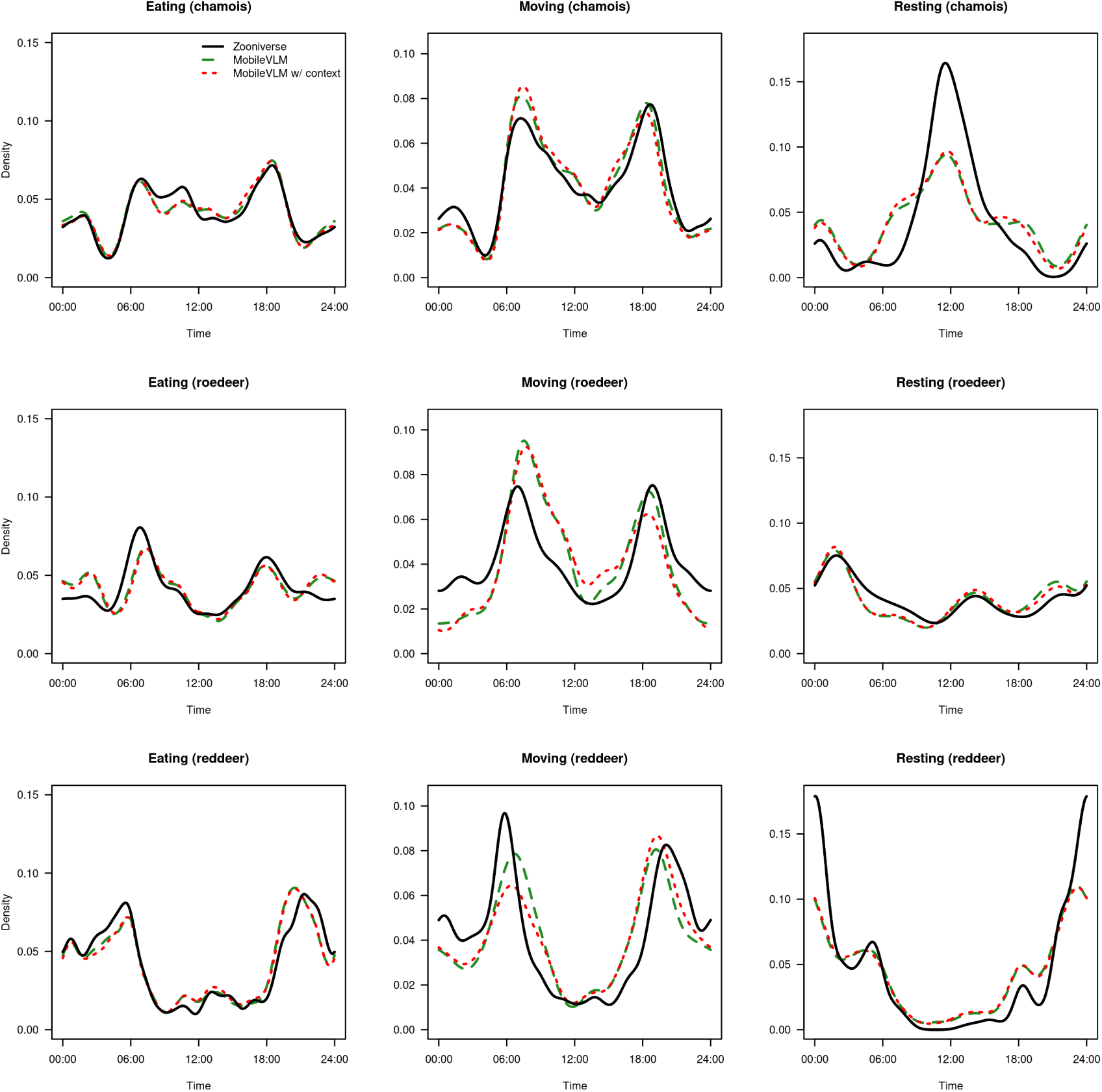
Activity patterns of MobileVLM V2 with and without day-night context, using the complete dataset.

**Figure 10:**
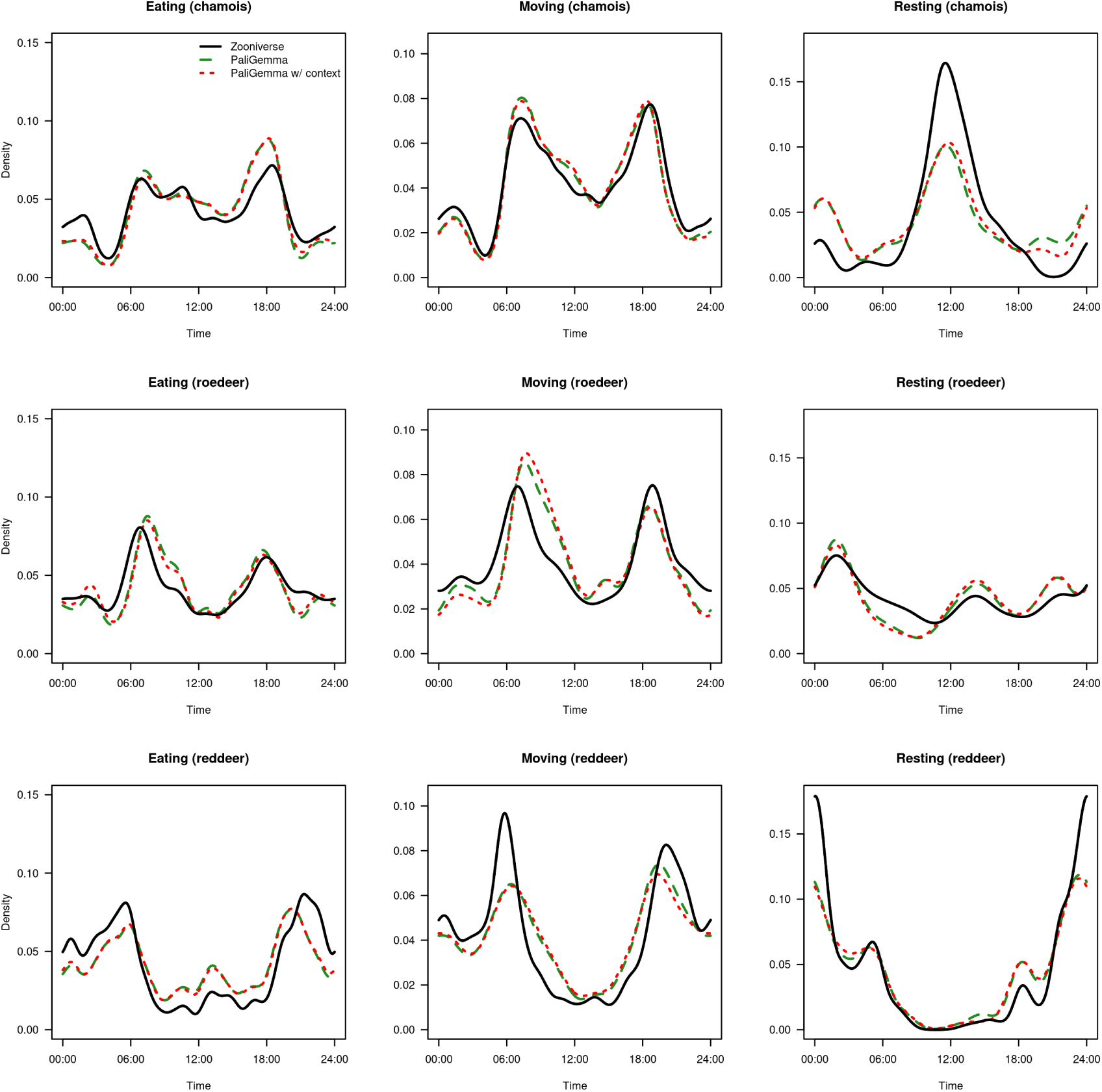
Activity patterns of PaliGemma, with and without day-night context, using the complete dataset.

**Table 5:**
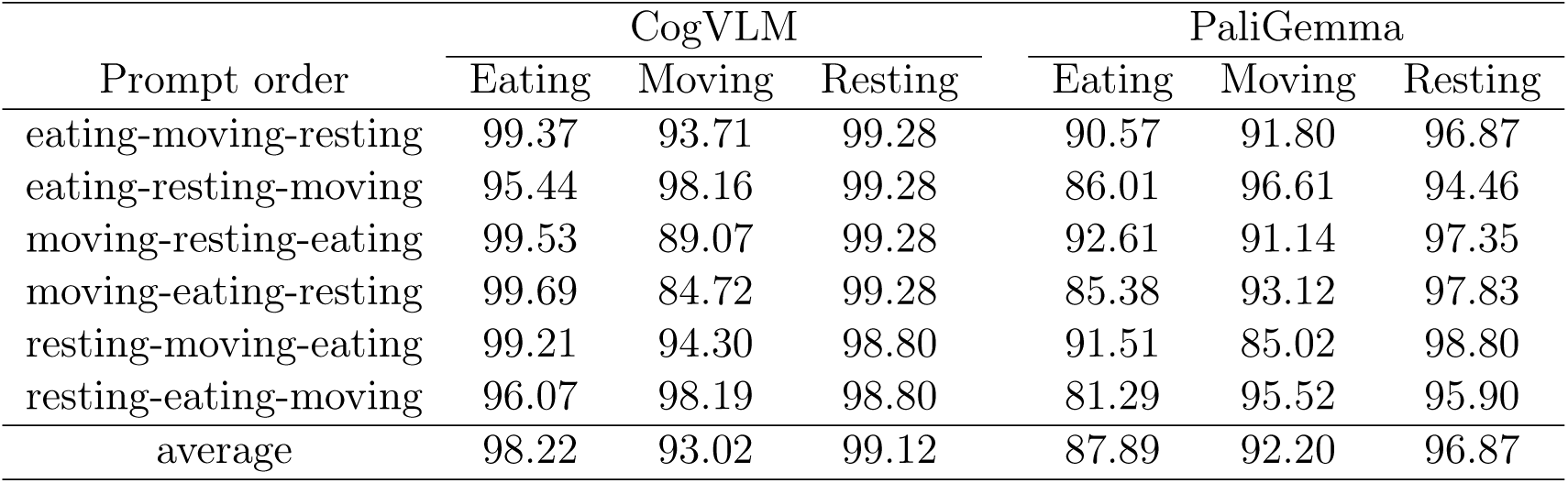
Behavior classification accuracy of the CogVLM and PaliGemma for different prompt permutations: in each prompt, the behaviors are given in a different order. For example, in the first row the prompt would be "Is the animal in the image eating, moving or resting?" and in the second row the prompt would be "Is the animal in the image eating, resting or moving?". To avoid complicating the presentation, the results in the study are given for the first permutation, as this is the one we used before studying the permutations.

